# Quantification of Cyclin-CDK dissociation constants in living cells using fluorescence cross-correlation spectroscopy with green and near-infrared fluorescent proteins

**DOI:** 10.1101/2024.11.14.623553

**Authors:** Aika Toyama, Yuhei Goto, Yuhei Yamauchi, Hironori Sugiyama, Yohei Kondo, Atsushi Mochizuki, Kazuhiro Aoki

## Abstract

The cell cycle is a highly coordinated process governed by cyclin-bound cyclin-dependent kinases (CDKs). While the interaction between cyclin and CDK are well-documented, the dissociation constants (K_d_) between specific cyclin-CDK pairs within living cells remain poorly understood. Fluorescence cross-correlation spectroscopy (FCCS) enables the quantification of the K_d_, but challenges remain in selecting an optimal pair of fluorescent molecules for FCCS in a living cell. In this study, we demonstrate that mNeonGreen and phycocyanobilin-bound miRFP670 represent a suitable pair for FCCS in living cells from the viewpoint of high photostability and low bleed-through. This fluorescent protein pair enables us to measure the K_d_ values of the cyclin-dependent kinase Cdc2 and B-type cyclin Cdc13 in fission yeast cells. Moreover, we conducted a comprehensive analysis of the K_d_ values for 36 cyclin-CDK complexes, formed by 9 distinct cyclins and 4 CDKs, in mammalian cells, including unconventional cyclin-CDK pairs. These findings provide insights into the redundancy of cyclin-CDK binding in cell cycle progression, with potential implications for understanding cell cycle regulation in both fission yeast and higher eukaryotes.

## Introduction

The cell cycle represents a highly coordinated process, by which eukaryotic cells divide into two daughter cells. The cell cycle comprises four phases, *i.e.*, G1, S, G2, and M phases, and these four cell cycle stages progress irreversibly in sequence. Cell cycle regulation is crucial for maintaining the homeostasis of eukaryotic cells and multicellular organisms. Altered cell cycle regulation is a hallmark of malignant tumors ^1^, and consequently the cell cycle is tightly controlled by multiple mechanisms.

Cyclin and cyclin-dependent kinases (CDKs) play an essential role in cell cycle regulation ^2^. Cyclin binds to and activates CDKs, thereby facilitating the cell cycle progression through the phosphorylation of CDK substrates. The first identification of a CDK was in the fission yeast *Schizosaccharomyces pombe*, where it was found in the *cdc2* gene ^3^. The *S. Pombe* genome encodes a single CDK gene, designated *cdc2*, and a single essential cyclin, *cdc13* (which is homologous to cyclin B). In fission yeast, it has been reported that the level of CDK activity mediated by the Cdc13-Cdc2 complex is a key determinant of the progression of each cell cycle phase ^4–7^. In contrast, mammalian cells predominantly utilize a variety of CDKs (CDK4, CDK6, CDK2, and CDK1) and cyclins (cyclin D, cyclin E, cyclin A, and cyclin B) for their cell cycle progression. The expression of cyclins is highly dependent on the cell cycle phase, whereas the expression level of CDKs remains constant ^8^. Accumulated evidence indicates that the formation of cell cycle phase-specific cyclin-CDK complexes ensures each cell cycle progression; cyclin D-CDK4/6 at G1 phase, cyclin E-CDK2 at G1/S phase ^9^, cyclin A-CDK2 at S-G2 phase ^10^, and cyclin B-CDK1 phase at G2/M phase ^11^. Meanwhile, it has been also reported that deficiencies in specific cyclins or CDKs are compensated for by other cyclins or CDKs. Mouse embryos in the early stages of development require only cyclin A2 and cyclin B1 for cell proliferation ^12,13^. Furthermore, CDK1 is the only essential CDK in mouse embryogenesis up to midgestation, and cyclin D1, cyclin D2, and cyclin E1 bind to CDK1 in embryos lacking all CDK2, CDK4, and CDK6 ^14^. These reports strongly implicate the redundancy in the function of cyclins and CDKs in cell cycle progression. However, although the interaction between cyclin and CDK are qualitatively well-documented, the quantitative understanding of binding affinity between specific cyclin-CDK pairs within living cells remains poorly understood.

The physicochemical interaction between proteins is of great importance for their functions, and the strength of protein-protein interaction is defined by the dissociation constant (K_d_). The K_d_ values have been determined through a series of *in vitro* experiments, including co-immunoprecipitation experiments, sedimentation equilibrium using analytical ultracentrifugation, surface plasmon resonance (SPR), and isothermal titration calorimetry (ITC). Fluorescence cross-correlation spectroscopy (FCCS) allows us to determine the K_d_ value of two molecules within living cells ^15^. In FCCS, fluctuations in the temporal profiles of two fluorescent species within the confocal volume (∼ 1 fL) are captured, followed by the cross-correlation analyses of time courses of the two fluorescence signals. The measurement of FCCS requires the continuous exposure of excitation light at the focal volume for a duration ranging from tens of seconds to minutes. However, due to the limited cell volume (∼ 5 pL in cultured cells ^16^ and 130 fL in fission yeast ^17^), prolonged and continuous exposure to excitation light may cause the photobleaching of the two fluorescent molecules. It is therefore essential that the two fluorescent molecules employed in FCCS should be tolerant to photobleaching. The green fluorescent protein monomeric enhanced green fluorescent protein (mEGFP) and mNeonGreen (mNG) have been demonstrated to exhibit robust resistance to photobleaching, thereby enabling their use in fluorescence correlation spectroscopy (FCS) measurements ^18,19^. On the other hand, the selection of a second fluorescent protein for FCCS was open to further consideration. However, the employment of the GFP-RFP pair for FCCS remains technically challenging due to their fast photobleaching and substantial triplet fraction of mCherry ^18^. HaloTag covalently attached to tetramethylrhodamine (TMR) provides a solution to these issues ^20^; however, the incubation of HaloTag ligands is required for the labeling of HaloTag proteins ^21^.

In this study, we report that a pair of mNG and miRFP670, a near-infrared fluorescent protein (iRFP), is a suitable pair for FCCS in living fission yeast and mammalian cells. iRFPs have been developed through the engineering of bacterial phytochromes ^22^. In contrast to conventional fluorescent proteins, such as GFP, iRFPs require a linear tetrapyrrole as a chromophore such as biliverdin IXa (BV) and phycocyanobilin (PCB). Among iRFPs, miRFP670 ^23^ is derived from the bacterial phytochromes of *Rhodopseudomonas palustris* RpBph1. By employing the pair of mNG and miRFP670, we succeeded in quantifying K_d_ values of cyclins and CDKs in fission yeast and mammalian cells using FCCS. The K_d_ values of all cyclin-CDK pairs measured in this study provide an insight into the roles played by not only canonical pairs but also non-canonical pairs in cell cycle progression.

## Materials and Methods

### Plasmids

All the plasmids used in this study are summarized in Table S1 with benchling links including their sequence and map information. The cDNA of human cyclin A1, cyclin A2, cyclin B1, cyclin B2, cyclin D1, cyclin D3, and cyclin E2 were amplified from cDNA synthesized from RNA extracted from HeLa cells by reverse transcription. The cDNA for cyclin D2 and cyclin E1 were synthesized by Fasmac. The cDNA of CDK1, CDK2, CDK4, and CDK6 were obtained from the Human Kinase OFR Kit from Hahn/Root Labs ^24^ (Addgene plasmids 23776, 23777, 23778, 23688, respectively). The cDNA of mNG and miRFP670 was inserted into the pCAGGS vector ^25^ to generate pCAGGS-mNG and pCAGGS-miRFP670 for negative control of FCCS experiments, respectively. For positive control of FCCS, the cDNA of miRFP670 was subcloned into pCAGGS-mNG to generate pCAGGS-mNG-miRFP670. The amino acid sequence (EAAAK)×3, a rigid linker ^26^, was inserted in between mNG and miRFP670 to minimize fluorescence resonance energy transfer (FRET) from mNG to miRFP670. To obtain the plasmids expressing cyclin fused with mNG, the cDNAs of cyclins were subcloned into the pCAGGS-mNG vector to obtain pCAGGS-cyclin A1-mNG, pCAGGS-cyclin A2-mNG, pCAGGS-cyclin B1-mNG, pCAGGS-cyclin B2-mNG, pCAGGS-cyclin D1-mNG, pCAGGS-cyclin D2-mNG, pCAGGS-cyclin D3-mNG, pCAGGS-cyclin E1-mNG, and pCAGGS-cyclin E2-mNG. The cDNA of CDKs was inserted into pCAGGS-miRFP670 to generate pCAGGS-CDK1-miRFP670, pCAGGS-CDK2-miRFP670, pCAGGS-CDK4-miRFP670, and pCAGGS-CDK6-miRFP670. For fission yeast endogenous tagging, miRFP670 was subcloned into a pFA6a vector using NEBuilder. To generate the fission yeast stable expression plasmid for miRFP670, miRFP670-mNeonGreen, mScarlet-I-mNeonGreen, and mCherry2-mNeonGreen, each gene was subcloned into pSKI vectors ^7,27^.

### Fission yeast strains and culture

All fission yeast strains used in this study are summarized in Table S2 along with their origins. Unless otherwise noted, the growth medium and other aspects of the experimental procedures for fission yeast were based on those described previously ^28^. The transformation protocol was adapted from a previously reported one ^29^.

### Photobleaching assay

A Leica SP8 Falcon confocal microscope (DMI8; Leica) was used for the photobleaching assay (Fig. 1). The microscope was equipped with an objective lens, the HC PL APO 63×/1.20 W motCORR CS2. Fission yeast cells expressing mScarlet-I-mNG, mCherry-2-mNG, or miRFP670-mNG under the control of the Padh1 promoter were mounted on standard glass slides and covered with a cover glass. The cells were illuminated using a white light pulsed laser with the following wavelengths; 488 nm for mNG, 561 nm for mScarlet-I and mCherry2, and 633 nm for miRFP670. The laser power was set to 1%, 3%, 5%, 10%, 20%, 40%, and 100% for mScarlet-I, mCherry, and miRFP, and 1% for mNG, which was utilized to select the cells expressing a comparable amount of fluorescent proteins. Leica HyD SMD detectors were used with the detection wavelength range of 490-540 nm for mNG, 640-795 nm for mScarlet-I and mCherry2, and 650-795 nm for miRFP670. The cytoplasm of the individual fission yeast cells was continuously illuminated for 15 seconds without laser scanning.

**Figure 1.**
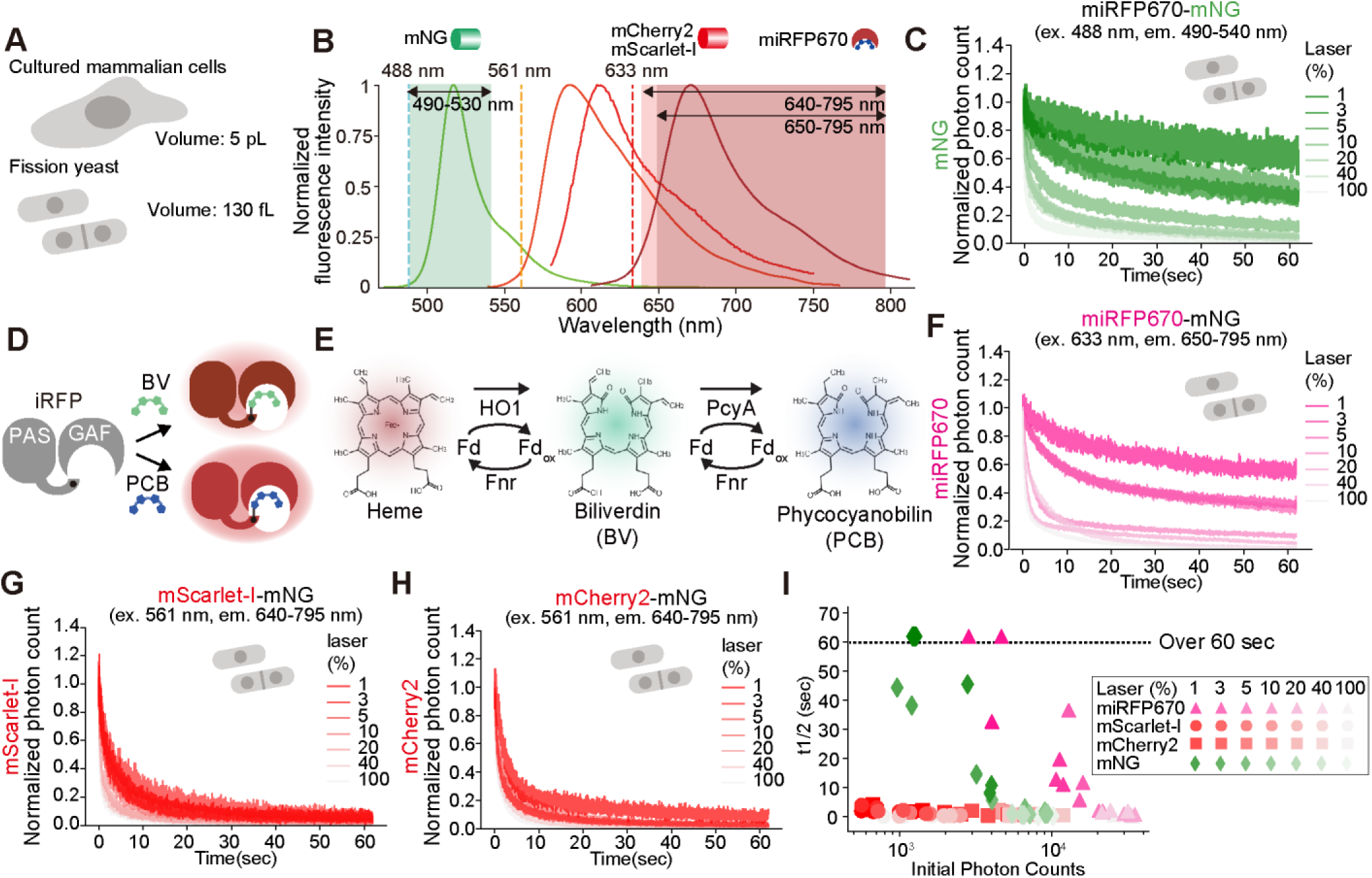
Comparison of photostability between RFPs and PCB-bound miRFP670. (A) Schematic illustration of a cultured mammalian cell and fission yeast with their cell volumes. (B) Fluorescence spectra of mNeonGreen (mNG, green), mCherry2 (orange), mScarlet-I (red), and miRFP670 (dark red). Dashed lines indicate excitation wavelengths for mNG (488 nm, light blue), mCherry2 and mScarlet-I (561 nm, yellow), and miRFP670 (633 nm, red). The emission channels were set to 490-530 nm (green), 640-795 nm (red), and 650-795 nm (dark red) for mNG, mScarlet-I and mCherry2, and miRFP670, respectively, as indicated by the pale bands. (C) Photobleaching assay for mNG in fission yeast cells by point-scan mode similar to FCS and FCCS, in which a 488 nm excitation laser (ex.) was continuously illuminated around the nucleus and photon counts at the emission wavelength of 490–540 nm (em.) were collected every 2 μsec for 60 sec. The photon counts recorded every 2 μsec were then normalized by dividing by the mean photon count value during the first 500 μsec. The normalized time series data were averaged (n = 3 cells), and the averaged data are plotted as a function of time. (D) Schematic of iRFP, which requires a linear tetrapyrrole such as biliverdin (BV) and phycocyanobilin (PCB) as a chromophore. (E) Schematic representation of the PCB biosynthesis pathway from heme. (F-H) Photobleaching assay for miRFP670 (F), mScarlet-I (G), and mCherry2 (H) in fission yeast cells by point-scan mode as in panel C. To reproduce the condition in FCCS, the excitation light for mNG (488 nm, 1%) was simultaneously exposed to the cells in addition to the excitation light for miRFP670 (633 nm, F), mScarlet-I (561 nm, G), or mCherry2 (561 nm, H). (I) The averaged half-life *t*_1/2_ (msec) of the indicated fluorescent proteins (n = 3) was plotted as a function of the raw values of their initial photon counts (the mean of the first 500 μsec).

### Culture and transfection of HeLa cells

HeLa cells were cultured in Dulbecco’s Modified Eagle’s Medium (DMEM) high glucose (Nacalai Tesque) supplemented with 10% fetal bovine serum (FBS; Sigma-Aldrich) at 37℃ in 5% CO2. HeLa cells lacking the *biliverdin reductase A* gene (*Blvra*) (HeLa/Blvra KO) ^30^ were utilized in FCCS experiments (Fig. 4). For FCCS measurement, HeLa/Blvra KO cells were seeded on a 35 mm glass base dish (IWAKI). One day after seeding, the cells were transfected with pCAGGS-SynPCB2.1 ^31^, pCAGGS-cyclin-mNG, and pCAGGS-CDK-miRFP670 (98:1:1 ratio) using 293fectin according to the manufacturer’s instructions. Two days after the transfection, the cells were subjected to FCCS experiments. HeLa/Blvra KO cells expressing SynPCB2.1 ^31^ (HeLa/Blvra KO/SynPCB) in a doxycycline-inducible manner were used in time-lapse imaging of either cyclins or CDKs (Fig. S3). For the time-lapse imaging of cyclins and CDKs during the cell cycle (Fig. S3), HeLa/Blvra KO/SynPCB cells were seeded on a 4 well glass base dish (Greiner), and treated with 1 µg/ml of doxycycline. One day after seeding, the cells were transfected with pCAGGS-cyclin-mNG or pCAGGS-CDK-miRFP670 using 293fectin. The cells were subjected to the time-lapse imaging 24 to 48 hours after transfection.

### Live-cell fluorescence imaging of HeLa cells

For the time-lapse imaging of either cyclins or CDKs (Fig. S3), an inverted fluorescence microscope (IX83; Olympus) were used, which was equipped with a Prime sCMOS camera (Photometrix, Tucson, AZ), and a dry objective lenses (UPLXAPO 20X, NA = 0.80, WD = 0.17 mm; Olympus). Filter settings were as follows: excitation filter, 475/28 nm (mNG), 633/22 nm (iRFP); emission filter, FF02-520/28 nm (mNG), BLP01-664R nm (iRFP). The microscope was controlled by MetaMorph software (ver. 7.10.4.407). During observation, cells were incubated in a stage incubator set to 37 °C and 5% CO2 (STXG-IX3WX, Tokai Hit). Images were taken every 10 minutes for 24 hours. Cells were tracked using a LIM tracker ^32^ to generate the montage images.

### FCS measurement in fission yeast cells

FCS data were obtained using a Leica SP8 Falcon confocal microscope (DMI8; Leica) equipped with an objective lens, a HC PL APO 63×/1.20 W motCORR CS2 objective. A white light pulsed laser was used to illuminate the samples. Fission yeast cells expressing mNG were measured at 488 nm (laser power 2%) excitation and 490–600 nm emission, and cells expressing miRFP670 were measured at 633 nm (laser power 0.75%) excitation and 640–795 nm emission. Time-series data of fluorescence fluctuations were obtained for 30 seconds. The output laser power at each wavelength is shown in the supplementary data (Fig. S2A). The structural parameter and effective confocal volume were calibrated using 1 μM Rhodamine 6G (TCI, R0039) in DDW based on the result that the diffusion constant of Rhodamine 6G in DDW is 414 µm^2^/s at room temperature ^20,33^.. The Rhodamine 6G solution was measured at 520 nm excitation and 550–700 nm emission.

### FCS and FCCS measurements in HeLa cells

FCS and FCCS data were obtained using a Leica SP8 Falcon confocal microscope (DMI8; Leica) equipped with an objective lens, HC PL APO 63×/1.20 W motCORR CS2. For HeLa cells, mNG was measured at 488 nm (laser power 10%) excitation and 490–600 nm emission, and cells expressing miRFP670 (laser power 10%) were measured at 633 nm excitation and 650–795 nm emission. During observation, cells were incubated in a stage incubator set at 37 °C. The structural parameter and effective confocal volume were calibrated as described above, and time-series data of fluorescence fluctuations were obtained for 15 seconds.

### Data analysis for FCCS

All data processing and analysis were performed using Python 3.12.4, with key libraries including NumPy for numerical operations, Pandas for data handling, Matplotlib for plotting, SciPy for signal processing and curve fitting, and the multipletau package for efficient computation of correlation functions.

The recorded fluorescence signals were used to calculate auto- and cross-correlation functions. The fluorescence time series data were first detrended and rescaled for photobleaching correction prior to subsequent analysis. We employed a detrending method based on a combination of moving average and Savitzky-Golay smoothing. For each fluorescence trace, a moving average with a window of 1500 data points was applied to remove high-frequency noise. The sparsely sampled moving-averaged trace was then smoothed using the Savitzky-Golay filter with a window length of 1501 points. This smoothed trace was then used to create an interpolation function that provided a continuous, smoothed baseline across the entire time series. Each original fluorescence trace was corrected by normalizing it by the square root of the ratio of the smoothed baseline to its initial value at time zero. This normalization helps maintain the overall shape of the fluorescence decay while correcting for slow photobleaching trends. The detrended traces were then resmoothed using the same moving average window to prepare them for correlation analysis.

FCS and FCCS analyses were performed on the detrended fluorescence traces. The auto-correlation function of each channel and the cross-correlation function between the two channels were computed using the multipletau algorithm, which is efficient for calculating correlation functions over logarithmically spaced time delays. The auto-correlation function and the cross-correlation function G(τ) for each channel were modeled using a function that accounts for diffusion dynamics:

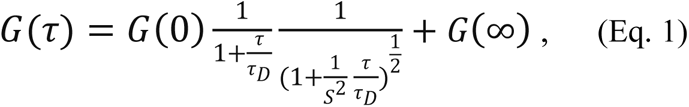

where *τ* is the lag time, *τ*_D_ is the diffusion time, 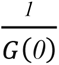 is the average number of fluorescent molecules in the observation volume, and *s* is the structure parameter. The correlation functions were fitted to these models using nonlinear least-squares optimization to extract the diffusion times and the average number of molecules. The fitting process was constrained by initial estimates and bounded by physically reasonable limits.

### Calculation of K_d_ for each cyclin-CDK pair

Using the parameters obtained from the analysis of the FCCS data, we calculated the dissociation constant K_d_ as described previously ^20^. Briefly, we obtained the number of molecules for green (*N*_G_), near-infrared (*N*_R_), and complexes (*N*_C_) by fitting for auto- and cross-correlation functions. Using the effective volume (*V*_eff_) calculated from the measurement and fitting of Rhodamine 6G, and Avogadro’s number (*N*_A_), `the concentration of each molecule can be expressed as follows:`

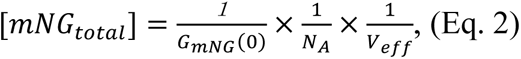

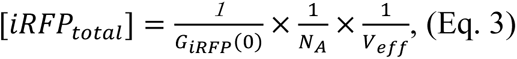

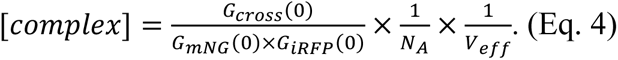

We also calculated the relative cross-correlation (RCC) value, which represents the fraction of molecular binding, as follows:

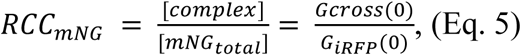

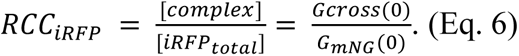

*RCC*_G_ and *RCC*_R_ are RCC of mNG and iRFP, respectively. These RCC values were further corrected by RCC values obtained the from positive control (*RCC*_PC_mNG_ and *RCC*_PC_iRFP_) and negative control (*RCC*_NC_mNG_ and *RCC*_NC_iRFP_) in order to quantitatively estimate to what extent the complex is formed ^20^. The corrected complex concentration, [*cComplex*], was given as follows:

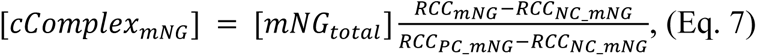

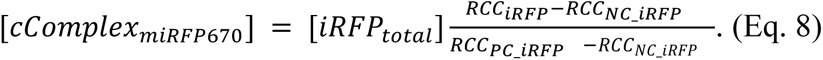

Finally, according to the definition of K_d_, the fraction of the complex in total mNG proteins or total iRFP proteins is derived as follows:

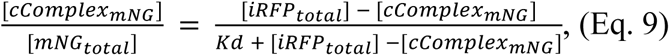

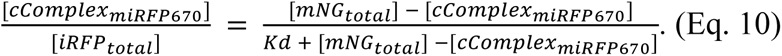

The fractions of the bound protein in total mNG and total iRFP (*i.e.*, the left-hand terms of equations 9 and 10) were plotted as a function of free (unbound) iRFP and mNG, respectively. The K_d_ value was obtained by nonlinearly fitting of the experimental data with equations 9 and 10 and by averaging of these fitted values. Python 3.12.4 with SciPy was used for curve fitting.

In FCCS in fission yeast, endogenous Cdc13 and Cdc2 are tagged with mNG and miRFP670, so there is little cell-to-cell variation in their expression levels. Therefore, instead of obtaining K_d_ values by nonlinear fitting, K_d_ was calculated using Equation 11 and 12, and average values were regarded as K_d_ values for each cell.

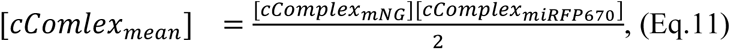

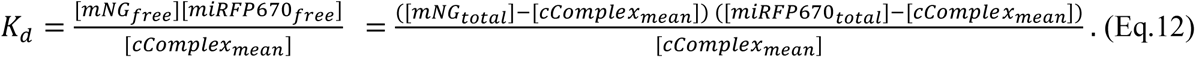

#### Images and data analysis

All fluorescence imaging data were analyzed and quantified using Fiji/ImageJ (https://fiji.sc/). Background was subtracted from all images using the rolling ball method.

## Results

### miRFP670, a near-infrared fluorescent protein, is suitable for simultaneous imaging with mNeonGreen

In order to perform FCCS in a reproducible manner, we are trying to find a better fluorescent protein pair that is bright, crosstalk free, and highly resistant to photobleaching. For this purpose, we used the fission yeast *Schizosaccharomyces pombe*, because the volume of a fission yeast cell (130 fL/cell) is about 40 times smaller than that of a mammalian cultured cell (5 pL) (Fig. 1A), making it much more sensitive to the effect of photobleaching. We first selected mNG as a green fluorescent protein for FCS and FCCS (Fig. 1B) because of its brightness and resistance to photobleaching ^18,19^. We overexpressed a fusion protein miRFP670-mNG in fission yeast cells, and investigated the photobleaching of mNG using point scan mode similar to FCS and FCCS, in which a 488 nm excitation laser was continuously illuminated at the cytoplasm and photon counts were collected every 2 μsec for 60 sec. As expected, mNG exhibited tolerance to the photobleaching when excited at low laser power (< 5%) (Fig. 1C).

Next, we investigated the possibility of iRFPs being applicable to FCS/FCCS with mNG, since the fluorescence spectra of iRFPs have little overlap with those of green fluorescent proteins (Fig. 1B). A linear tetrapyrrole chromophore, such as biliverdin IXa (BV) and phycocyanobilin (PCB), is required for the fluorescence of iRFPs. Although BV is biosynthesized from heme by heme oxygenase-1 (HO-1) in a wide range of living organisms (Fig. 1E), the fission yeast *S. Pombe* has lost an HO-1 gene during evolution ^27^. In cyanobacteria, PCBs are mainly generated from BV by phycocyanobilin: ferredoxin oxidoreductase (PcyA) ^34^(Fig. 1E). These tetrapyrroles are covalently attached to iRFPs and are therefore capable of emitting infrared fluorescence. We have previously reported that PCB-bound iRFPs exhibit higher fluorescence intensity than BV-bound iRFPs ^27^ (Fig. 1D). In addition, we have successfully developed the system for efficient PCB biosynthesis, called SynPCB, in mammalian cells ^30,31^, fission yeast ^27^, and *C. elegans* ^35^.

Among iRFPs, we chose miRFP670 because of the higher brightness compared to other iRFPs ^27^. The fission yeast cells expressing SynPCB2.1 and miRFP670-mNG proteins were exposed to the 633 nm excitation laser in the point-scan mode as shown in Figure 1C. To reproduce the condition in FCCS, the excitation light for mNG (488 nm, 1%) was simultaneously exposed to the cells. PCB-bound miRFP670 fluorescence decayed slowly, but retained more than 50% of its initial fluorescence intensity after 10 seconds at laser intensities of 1% and 3% (Fig. 1F). On the other hand, mScarlet-I and mCherry2, which have been reported to be resistant to photobleaching ^36^, were rapidly photobleached in the fission yeast cells even at the low laser power of 561 nm excitation laser (Fig. 1G and 1H). To compare the fluorescence intensity and photobleaching characteristics, we quantified the half-life *t*_1/2_, the time at which the fluorescence intensity is reduced to a half of the initial one, from each time-series data of mNG, miRFP670, mScaler-I, and mCherry, and plotted it as a function of the initial photon count (the mean photon count of first 500 μsec) (Fig. 1I). miRFP670 and mNG showed higher or comparable initial photons count than mScarlet-I and mCherry2, even under laser conditions where the half-life *t*_1/2_ exceeds 60 seconds (Fig. 1I). The reason why the initial photon counts of mScarlet-I and mCherry2 were much lower than that of miRFP670 was that the fluorescence detection channel of mScarlet-I and mCherry2 (640-795 nm) was shifted from their optimal emission wavelength (around 600 nm) in order to reduce the bleedthrough of mNG fluorescence (Fig. 1B). We confirmed the linear relationship between the laser power settings of the microscope (%) and light intensity focused through the objective lens at 488 nm, 561 nm, and 633 nm lasers (Fig. S1). These results indicate that PCB-bound miRFP670 is much more suitable for FCS and FCCS analysis with mNG than mScarlet-I and mCherry2, in terms of fluorescence intensity and photostability.

### FCS and FCCS by using mNG and miRFP670

We investigated whether miRFP670 can be used for FCS and FCCS analysis in fission yeast cells. In FCS, a target protein A fused to miRFP670 is expressed in cells and continuously excited by an excitation laser (Fig. 2A, left) to obtain the time-series data of fluorescence fluctuation in a tiny confocal volume (approximately 1 fL) with a confocal laser scanning microscope (Fig. 2A, middle). Fluctuation in fluorescence intensity is attributed to the random process of the target proteins entering and exiting the confocal volume. Using time-series data, an auto-correlation function (ACF) is calculated, and the average number of fluorescent proteins in the confocal volume is obtained from the y-intercept value of the ACF curve (Fig. 2A, right); the y-intercept value is inversely proportional to the average number of molecules (see Materials and Methods for more details). In FCCS, two target proteins A and B, labeled with mNG and miRFP670, respectively, are expressed in cells (Fig. 2B, left). As in FCS, fluorescence intensity fluctuations of mNG and miRFP670 are measured and their ACFs are calculated to obtain the average number of molecules in the confocal volume (Fig. 2B, middle and right). The two time-series data of mNG and miRFP670 are further analyzed to calculate the cross-correlation function (CCF) (Fig. 2B, right). The y-intercept values of CCF and ACF allow us to estimate the average number of molecules forming the complex between protein A and B (see Materials and Methods for more details).

**Figure 2.**
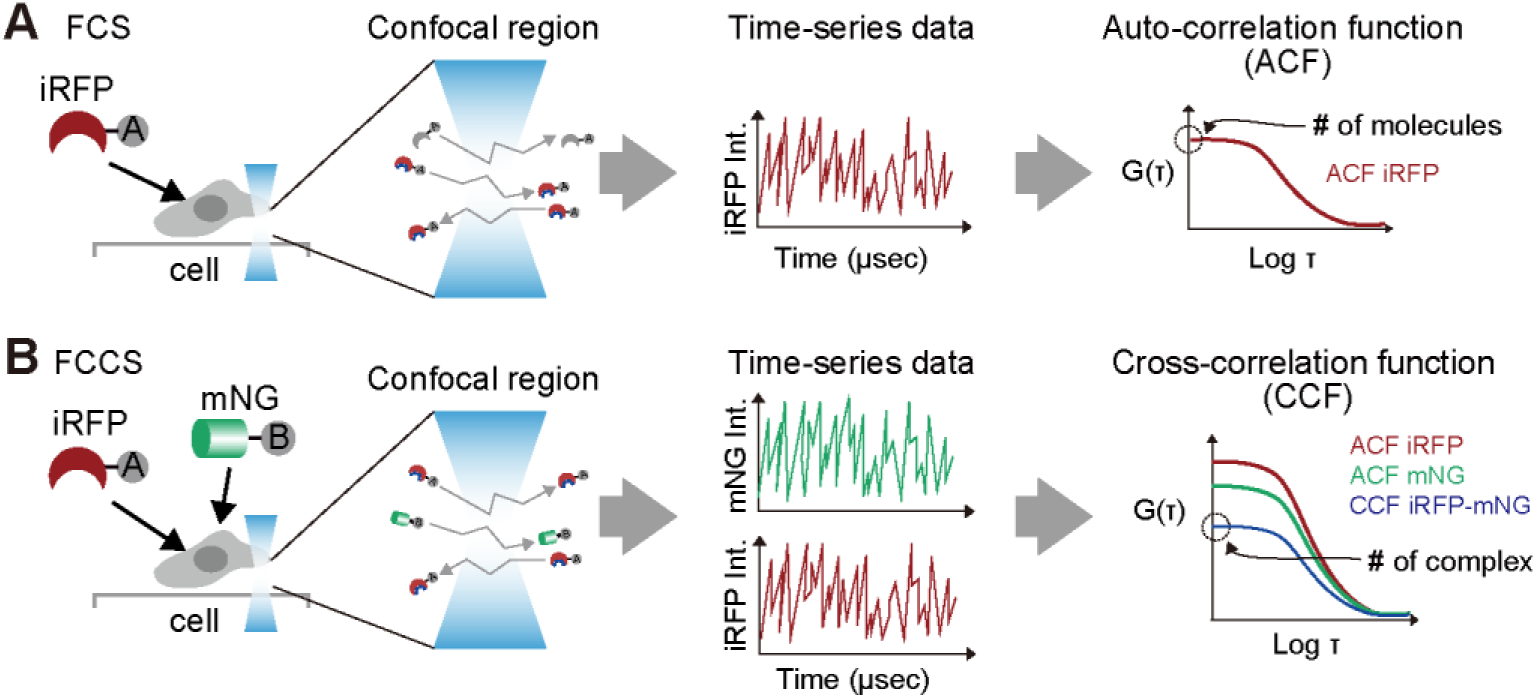
Schematic representation of FCS and FCCS with mNG and miRFP670. (A and B) Experimental procedure of measurement and analysis of FCS (A) and FCCS (B) by using mNG and miRFP670. In FCS (A), protein A fused with miRFP670 and SynPCB are expressed in cells, and time-series of PCB-bound miRFP670 fluorescence is recorded in point-scan mode with a confocal microscope. The auto-correlation function (ACF) is calculated and plotted as a function of the logarithm of the lag time, τ. In the ACF, the inverse of the *y*-intercepts represents the mean number of fluorescent molecules in the confocal volume (∼ 1 fL). In FCCS (B), proteinA and B fused to miRFP670 and mNG, respectively, and SynPCB are expressed in cells. From the time series data of miRFP670 and mNG, the ACFs for miRFP670 (red curve) and mNG (green curve) and the cross-correlation function (CCF) (blue curve) can be calculated. The mean number of complexes is calculated from the y-intercept values of ACF and CCF.

### Quantification of the dissociation constant of the Cdc13-Cdc2 complex in a living fission yeast cell

Next, we tested miRFP670 for FCS and FCCS in fission yeast cells. To employ miRFP670 in fission yeast, we used a fission yeast strain that expressed SynPCB2.1 under the control of the *adh* promoter ^27,31^. We used a fusion protein of mNG and miRFP670, *i.e.*, mNG-miRFP670, as a positive control for FCCS. As expected, the mNG-miRFP670 showed substantial amplitude of the cross-correlation curve (Fig. 3A). On the other hand, the fission yeast strain expressing mNG and miRFP670 separately, as a negative control, exhibited almost no amplitude of the cross-correlation curve (Fig. 3B). Based on these results, we calculated the relative cross-correlation (RCC) values, which indicate the proportion of fluorescent molecules that form a complex. The RCC values are approximately 0.7 and 0.1 for the positive and negative controls, respectively (Fig. 3C). The reason why the RCC in the positive control was less than 1 could be due to the presence of fusion proteins in which either mNG or miRFP did not fluoresce, the mismatch of confocal volumes at the different excitation wavelengths, and/or the measurement noise of the detection channels. In subsequent experiments, these values were used to correct for complex concentration (for more details, refer to the Material and Methods).

**Figure 3.**
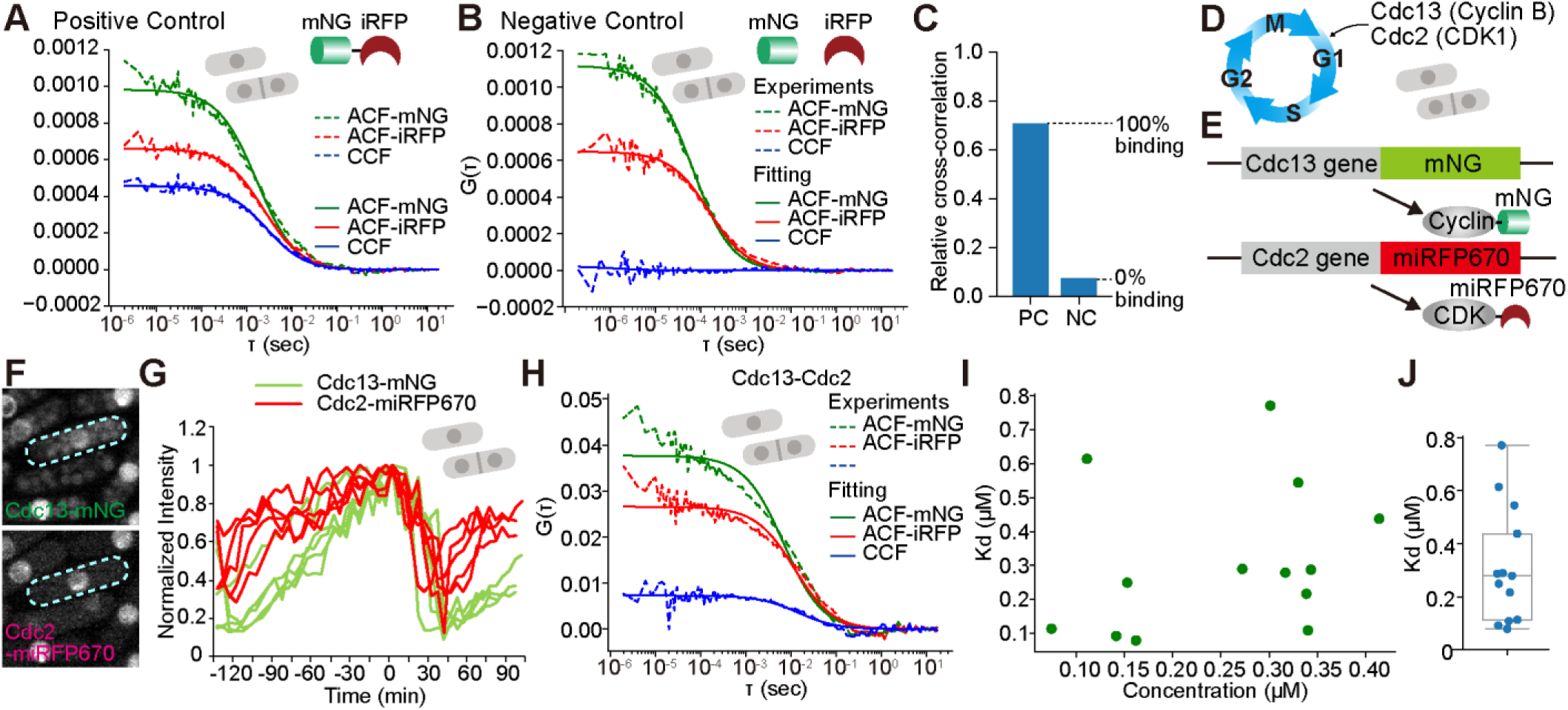
Quantification of K_d_ values between Cdc13 and Cdc2 by FCS and FCCS in fission yeast cells. (A and B) Representative ACF and CCF of the positive control (PC) with mNG-miRFP670 fusion protein (A) and the negative control (NC) with mNG and miRFP (B). The dashed and solid lines represent the ACF/CCF functions obtained from the experimental time-series data and the fitted curves, respectively. (C) Relative cross-correlation (RCC) for PC and NC. These values were used to correct the concentration of complexes. The RCC values for PC and NC were regarded as 100% binding and 0% binding, respectively. (D) Schematic illustration of the cell cycle of fission yeast. (E) The fission yeast strain endogenously expresses Cdc13-mNG and Cdc2-miRFP670. (F) Subcellular localization of Cdc13-mNG and Cdc2-miRFP670 in fission yeast cells. (G) Time-course of Cdc13-mNG and Cdc2-miRFP670 mean nuclear fluorescence intensity throughout the cell cycle. Light green and red lines indicate Cdc13-mNG and Cdc2-miRFP670, respectively. The time-series data were aligned with the maximum of Cdc13-mNG as t = 0. (H) Representative auto- and cross-correlation curves of Cdc13-mNG and Cdc2-miRFP670. (I) The K_d_ values for each of 13 cells plotted as a function of Cdc13-mNG fluorescence intensity. (J) The box plot of the K_d_ values with the whiskers donating 1.5 times the interquartile range. The box extends from the first to the third quartile. The blue dot indicates data from individual cells.

It is widely accepted that the cell cycle of fission yeast is regulated by a single CDK, Cdc2, and four cyclins, Cdc13, Cig1, Cig2, and Puc1 ^37^. Among the cyclins, only Cdc13 is an essential cyclin, and thus only a Cdc13-Cdc2 pair is sufficient for the cell cycle progression in fission yeast ^5^(Fig. 3D). Therefore, we quantified the dissociation constant of the endogenous Cdc13 and Cdc2 complex in a proliferating fission yeast cell. First, an *mNG* and *miRFP670* gene were integrated into at the the 3’ end of the *cdc13* and *cdc2* gene, respectively (Fig. 3E). The time-courses of Cdc13-mNG and Cdc2-miRFP670 in the nucleus during cell cycle progression were quantified, and Cdc13-mNG and Cdc2-miRFP670 were mainly localized at the nucleus as had been previously reported ^38^(Fig. 3F). Moreover, consistent with the previous finding ^38^, the fluorescent intensity of Cdc13-mNG gradually increased during the G1, S, and G2 phases, followed by a rapid decline at the metaphase of mitosis (Fig. 3G, green lines). Consequently, the Cdc13-mNG fluorescence intensity allows us to infer the cell cycle stage. The fluorescent intensity of Cdc2-miRFP670 at the nucleus slightly increased during G1, S, and G2 phases and disappeared at the metaphase (Fig. 3G, red lines). Finally, we determined the concentration of Cdc13-mNG, Cdc2-miRFP670, and their complex at the nucleus by using FCS and FCCS (Fig. 3H), and we estimated the K_d_ values in each cell (Fig. 3I). The cells expressing varying amounts of Cdc13-mNG were subjected to the FCS and FCCS analysis for obtaining the K_d_ values at different phases of the cell cycle. We found only a weak positive correlation (*r* = 0.46) between cell cycle stage (i.e., Cdc13-mNG concentration) and the K_d_ of Cdc13-mNG and Cdc2-miRFP670 complex (Fig. 3I), suggesting that Cdc2 interacts with Cdc13 independently of cell cycle phases-specific modification or regulation in fission yeast cells. The mean K_d_ value calculated for each of the 13 cells was 0.31±0.22 μM.

### Quantification of dissociation constants of possible pairs between 9 types of cyclins and 4 types of CDKs in a living HeLa cell

Next, we applied FCS and FCCS with the mNG and miRFP670 pair to the comprehensive analysis of the interaction between cyclins and CDKs in mammalian cells. We chose 9 types of cyclins (cyclin D1, D2, D3, E1, E2, A1, A2, B1, and B2) and 4 types of CDKs (CDK4, CDK6, CDK2, and CDK1). Note that cyclin B3 gene could not be cloned from the cDNA library of HeLa cells.

To demonstrate whether FCS and FCCS analysis using mNG and miRFP670 is feasible in mammalian cells, we did positive and negative control experiments as in Figure 3A and 3B. Notably, to label miRFP670 with PCB, we used biliverdin reductase A gene (*Blvra*) KO HeLa cells and transiently introduced plasmids expressing SynPCB2.1 for PCB synthesis ^31^. The depletion of Blvra, which catalyzes the conversion of BV and the PCB to bilirubin and phycocyanorubin, respectively, results in an increase in the amount of BV and PCB ^30^. The expression of SynPCB2.1 in parental HeLa cells increased miRFP670 fluorescence intensity by about 4-fold (Fig. S2A), because of the brighter chromophore, PCB, than BV. The depletion of Blvra also showed about 4-fold increases in miRFP670 fluorescence (Fig. S1A), resulting from the accumulation of BV. The combination of the SynPCB2.1 expression with the Blvra depletion further enhanced miRFP670 fluorescence by up to approximately 10-fold (Fig. S2A).The cells expressing a chimeric fusion protein of mNG-(EAAAK)*3-miRFP670 (mNG-miRFP670) were employed as the positive control, while the cells separately expressing mNG and miRFP670 served as the negative control (Fig. 4A and 4B). As anticipated, the positive control demonstrated a considerable y-intercept value for CCF, while the negative control did not (Fig. 4A and 4B, blue lines). IIt is noteworthy that the y-intercept values of the ACFs of mNG and miRFP670 in the positive control experiment (Fig. 4A) consistently indicated a greater number of miRFP670 than mNG (Fig. S2B). Given that mNG is known to be matured efficiently in living cells ^39^, it is reasonable to conclude that the majority of the miRFP670 proteins are the holo-form bound to PCB in Blvra KO HeLa expressing SynPCB2.1. From these results, the RCC of the positive control and the negative control was determined to estimate the dissociation constant (Fig. 4C). In our FCCS setup, the RCC values were within the range of 0.3 to 0.6 and 0.002 to 0.07 for the positive and negative controls, respectively (Fig. 4C).

**Figure 4.**
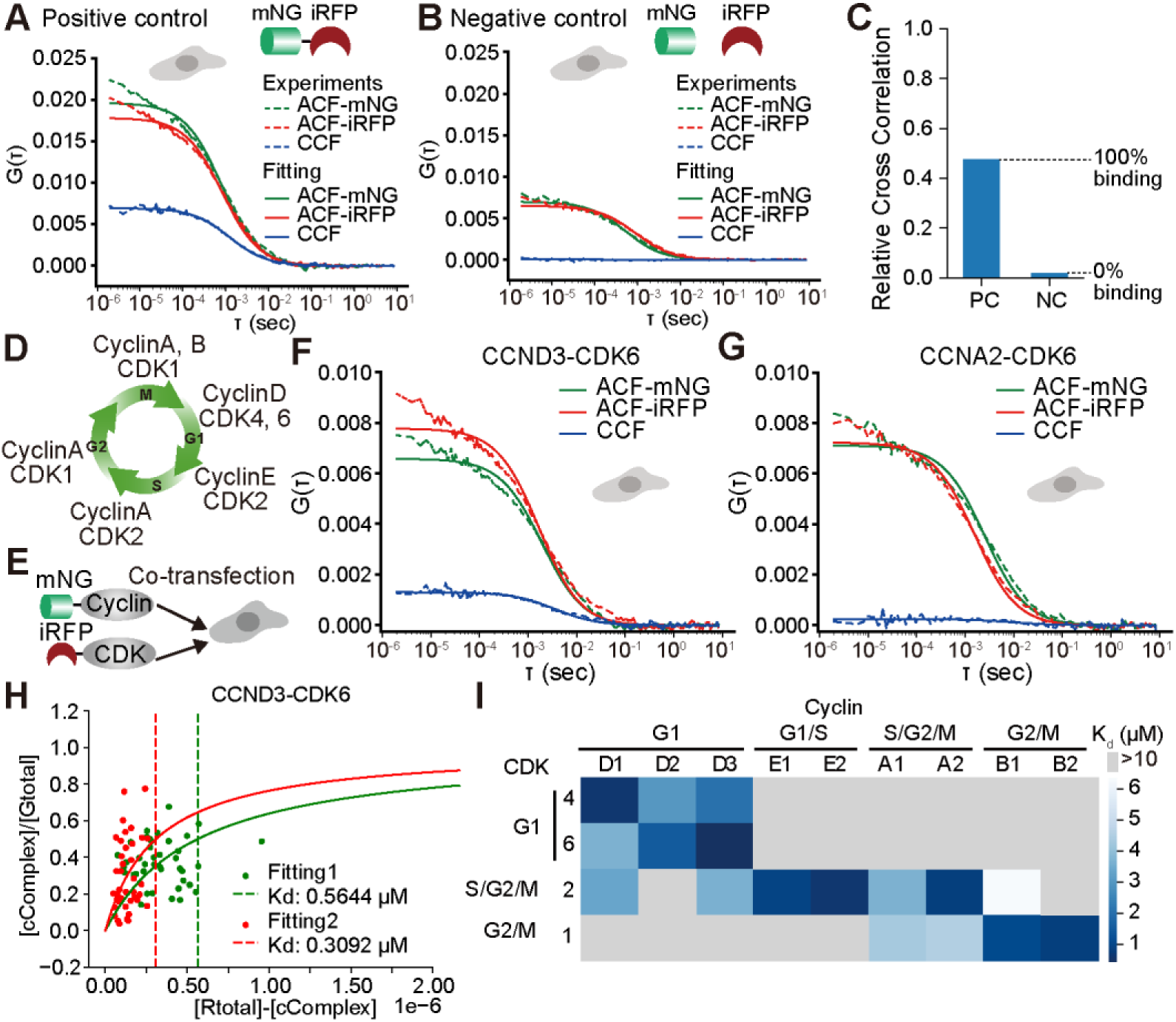
Quantification of K_d_ values between cyclins and CDKs by FCS and FCCS in HeLa cells. (A and B) Representative ACF and CCF of the positive control (PC) with mNG-miRFP670 fusion protein (A) and the negative control (NC) with mNG and miRFP (B). The dashed and solid lines represent the ACF/CCF functions obtained from the experimental time-series data and the fitted curves, respectively. For PC, *Blvra* KO HeLa cells were transfected with the plasmid encoding a fusion protein of mNG, the (EAAAK)×3 linker, and miRFP670 (mNG-miRFP670), and SynPCB. For NC, cells were transfected with the plasmid expressing SynPCB and miRFP670 and mNG separately, and SynPCB. Representative data are shown. (C) RCC for PC and NC in *Blvra* KO HeLa cells. These values were used to correct for the concentration of the complexes. (D) Schematic illustration of the mammalian cell cycle and the conventional cyclin-CDK pairs for each phase. (E) Schematic illustration of the construct used in the experiments. Cyclins and CDKs were labeled with mNG and miRFP670, respectively. *Blvra* KO HeLa cells were co-transfected with plasmids expressing cyclin-mNG, CDK-miRFP670, and SynPCB. (F and G) Representative auto- and cross-correlation curve of cyclin D3-CDK6 complex (F) and cyclin A2-CDK6 complex (G). (H) The data with the fitted curve for calculating the K_d_ of cyclin D3-CDK6 complex. (I) Heat map showing the K_d_ values between a possible pair of cyclins and CDKs. Gray tiles indicate that the K_d_ exceeds 10 uM.

Finally, we quantified the dissociation constants of 36 pairs of 9 cyclins and 4 CDKs complexes in HeLa cells. In the mammalian cell cycle, it is widely recognized that different pairs of cyclin-CDK complexes are formed to regulate the different stages of cell cycle progression. (Fig. 4D). In this analysis, we transiently overexpressed SynPCB, an mNG-fused cyclin, and an miRFP670-fused CDK in *Blvra* KO HeLa cells, and measured the concentrations of mNG-cyclin, miRFP670-CDK, and cyclin-mNG and CDK-miRFP670 complexes (Fig. 4E). For example, the cyclin D3-CDK6 pair exhibited a substantial y-intercept value of CCF (Fig. 4F), indicating the complex formation between cyclin D3 and CDK6. Meanwhile, the y-intercept value of the cyclin A2-CDK6 pair was almost zero (Fig. 4G), and therefore no interaction was detected between cyclin A2 and CDK6. Data were obtained from approximately 40 cells with varying levels of cyclin and CDK expression due to transient expression by lipofection. These data were plotted according to the equations 9 and 10 described in the Materials and Methods, and K_d_ values were obtained by fitting. The data with the fitted curve for cyclin D3-CDK6 are shown as an example (Fig. 4H), and that of the other pairs are shown in Fig. S4. The K_d_ values for all 36 pairs of cyclin-CDK complexes are represented as a heatmap (Fig. 4I). The K_d_ values of low affinity cyclin-CDK pairs that exceed 10 μM are assigned as gray tiles. As expected, the conventional cyclin-CDK pairs, *i.e.*, cyclin D-CDK4/6, cyclin E-CDK2, cyclin A-CDK2, cyclin A-CDK1, and cyclin B-CDK1 (Fig. 4D), showed relatively low K_d_ values (Fig. 4I). We found the difference between subtypes of cyclin D and cyclin A; cyclin D1 preferentially binds to CDK4, while cyclin D2 and D3 interact with CDK6 more strongly. Similarly, both cyclin A1 and A2 bind to CDK2 and CDK1, but the K_d_ value of the cyclin A2-CDK2 complex is much smaller than that of the others. Interestingly, the bindings of cyclin D1 and D3 to CDK2 and cyclin B1 to CDK2 were also observed, suggesting that unconventional cyclin-CDK pairs form in living cells. Furthermore, the K_d_ values of cyclin E1, E2 and cyclin A2 to CDK2 were comparable, indicating the redundancy in the role of cyclins in CDK2 activation. Cyclin E1 and E2 formed a complex with only CDK2.

## Discussion

In this study, we demonstrated that PCB-bound miRFP670 is much more suitable for FCS and FCCS analysis with mNG than mScarlet-I and mCherry2, in terms of fluorescence intensity and photostability. Under the optimal conditions, mScarlet-I and mCherry2 are brighter fluorescent proteins than miRFP670 ^40^. However, under the conditions of FCCS with mNG, it is necessary to shift the detection channel of red fluorescent proteins to longer wavelengths to reduce the bleedthrough of mNG fluorescence, which forces the use of stronger excitation light to obtain a sufficient number of photons. In addition, during FCCS, the mNG excitation light must also be irradiated at the same time. These two factors would accelerate the photobleaching of mScarlet-I and mCherry2 (Fig. 1). HaloTag attached to tetramethylrhodamine (TMR) allows FCCS measurement in cultured cells, because TMR is bright and resistant to photobleaching ^20,41^. However, it requires the addition of HaloTag-ligand and subsequent washout, and HaloTag-ligand does not enter in fission yeast uniformly. On the other hand, miRFP670 can be excited and detected under optimal conditions during FCCS with mNG. In addition, there is only a little absorption of miRFP670 at 488 nm, and therefore photobleaching by the 488 nm excitation light may be negligible. In principle, fluorophores that emit at longer wavelengths typically have lower energy excited states; this lower energy correlates with a reduced probability of intersystem crossing to the triplet state and a lower probability of photobleaching ^42^. For these reasons, miRFP670 is a better choice for FCCS with mNG.

We succeeded in measuring K_d_ values of cyclins and CDKs using FCCS in fission yeast and HeLa cells. To the best of our knowledge, many reports have demonstrated the cyclin-CDK binding detected by immunoprecipitation, but there are very few papers that have measured K_d_ values. Jonathon Pines and colleagues reported pioneering results using FCS and FCCS to measure dissociation constants after endogenous labeling of cyclin B1 and CDK1 with genome editing in RPE-1/hTERT cells ^43^. They arrested the cell cycle in the G1 phase by treating the cells with a CDK4/6 inhibitor, followed by the release of the G1 cell cycle arrest to synchronize the cell cycle. Then, they measure the dissociation constant in each cell cycle phase; their dissociation constants of cyclin B1 and CDK1 are 270 nM, 158 nM, and 112 nM after 6, 9, and 12 hours of cell cycle release, respectively. On the other hand, our present study showed the K_d_ value of 1.1 μM for cyclin B1 and CDK1 in HeLa cells (Fig. 4), which was almost an order of magnitude higher than the previous report. This discrepancy can be partly attributed to the overexpression of cyclins and CDKs, in which the endogenous (non-fluorescently labeled) molecules act as competitors, thereby overestimating the dissociation constants ^20,41^. Consistent with this, we measured the dissociation constants of endogenous cdc13/cyclin B and cdc2/CDK1 in fission yeast by using FCCS, and found that the dissociation constant of fission yeast cdc13 and cdc2 was 310 nM (Fig. 3), which is comparable to the dissociation constants of endogenous cyclin B1 and CDK1 in RPE-1/hTERT cells.

We comprehensively measured the dissociation constants of 36 different pairs of cyclins and CDKs, and, as expected, we detected the binding of conventional cyclin-CDK pairs, namely cyclin D-CDK4/6, cyclin E-CDK2, cyclin A-CDK2, cyclin A-CDK1, and cyclin B-CDK1 (Figure 4). As mentioned above, these K_d_ values would be overestimated, but to the best of our knowledge, these data demonstrate for the first time the direct comparison of the binding strength between cyclins and CDKs, which has been evaluated in the same cell type with the same experimental system. Interestingly, we unexpectedly recognized a weak binding between the unconventional cyclin-CDK pair, *i.e.*, cyclin D1/3-CDK2 and cyclin B1-CDK2. Previous studies have reported that a certain cyclin binds to unconventional CDKs and inhibits their function. For instance, several research groups have shown the presence of the cyclin D-CDK2 complex ^44^ and its inhibitory role ^45^ in CDK2 activity. Cyclin A-CDK1 complex also suppresses the expression of Cdc25 to restrict the G2/M phase progression ^46^. These results indicate the possibility that the unconventional cyclin-CDK complex exerts an inhibitory effect on cell cycle progression, thereby ensuring timely cell cycle progression. On the other hand, some reports support the idea that the interaction of cyclin with the unconventional CDK leads to the activation of CDK such as the cyclin D-CDK2 complex ^47,48^. When CRL4^AMBRA1^, which ubiquitinates cyclin D, is depleted, cyclin D levels increase, resulting in the accumulation of the cyclin D1-CDK2 and cyclin D3-CDK2 complex and the decrease in sensitivity to CDK4/6 inhibitors ^49,50^. Furthermore, the existence of an unconventional cyclin B1-CDK2 complex has been reported in CDK1 knock-out (KO) or knock-down (KD) cells ^51,52^. Although CDK1 depletion leads to cell cycle arrest, CDK2 overexpression rescues the cell cycle arrest by CDK1 depletion ^52^, suggesting the role of the unconventional cyclin B1-CDK2 in the cell cycle progression. The roles of such unconventional cyclin-CDK pairs remain to be elucidated, and the measurement of the intracellular K_d_ values allows us to quantitatively interpret when and to what extent these unconventional cyclin-CDK complexes are present and functioning during cell cycle progression.

Finally, we should mention the technical challenges of this study. The first challenge is the potential toxicity associated with the expression of SynPCB. It is necessary to express SynPCB in order to use PCB-bound iRFP. Although the cultured mammalian cells and fission yeast showed no toxicity, the expression of SynPCB caused toxicity in *Caenorhabditis elegans* ^35^. Thus, we should keep in mind the potential toxicity induced by the SynPCB expression. Second, the depletion of *Blvra* proteins is essential for achieving maximal labeling of iRFP (Fig. S2A) ^27,53^, which is preferable for FCCS analysis. Fission yeast and *C. elegans* do not acquire the *Blvra* gene, and therefore the SynPCB expression suffices to fully label iRFP by PCB ^27,35^. Third, we measured the dissociation constants between cyclins and CDKs using a transient overexpression system without cell cycle synchronization in HeLa cells (Fig. 4). Therefore, the K_d_ values were overestimated and averaged across cell cycle phases. It would be ideal to label endogenous cyclin and CDK using genome editing, synchronize the cell cycle, and measure the dissociation constant ^43^. Future studies should address these challenges to improve the accuracy of FCS and FCCS measurements in a variety of living organisms. This would allow for a deeper investigation and understanding of cyclin-CDK interactions.

## Abbreviations

BV: Biliverdin
PCB: Phycocyanobilin
FCS: Fluorescence Correlation Spectroscopy
FCCS: Fluorescence Cross-Correlation Spectroscopy
CDK: cyclin dependent kinase

## Acknowledgements

We thank all members of the Aoki Laboratory for their helpful discussions and assistance. Some fission yeast strains were provided by the National Bio-Resource Project (NBRP), Japan. This work was supported by the Live Imaging Center, Kyoto University.

## Funding

K.A. was supported by JSPS KAKENHI grants (nos. JP19H05798, JP22H02625, JP24H01416, and JP24K21981), NAGASE Science Technology Foundation, Takeda Science Foundation, the grant of Joint Research by the National Institutes of Natural Sciences (NINS)(NINS program No. OML012404), Joint Research of the Exploratory Research Center on Life and Living Systems (ExCELLS)(ExCELLS program No. 23EXC601), and the Cooperative Research Program (Joint Usage/Research Center program) of Institute for Life and Medical Sciences, Kyoto University. Y.G. was supported by a JST, ACT-X grant (no. JPMJAX22B8), and JSPS KAKENHI grants (nos.19K16050 and 22K15110).

## Author Contributions

Y.G., and K.A. designed the research; A.T., Y.Y., and Y.G. performed the experiments; A.T., Y.Y., Y. K., and Y.G. analyzed the data; A.T., Y.G., A.M., and K.A. wrote the manuscript; all authors approved the final manuscript.

## Competing Interests

The authors have no competing interests to disclose.

## Notes

### Competing Interest Statement

The authors have declared no competing interest.

