## Supplementary Information for "Quantification of Cyclin-CDK dissociation constants in living cells using fluorescence cross-correlation spectroscopy with green and near-infrared fluorescent proteins"

### Title:

**Supplementary Figure 1**

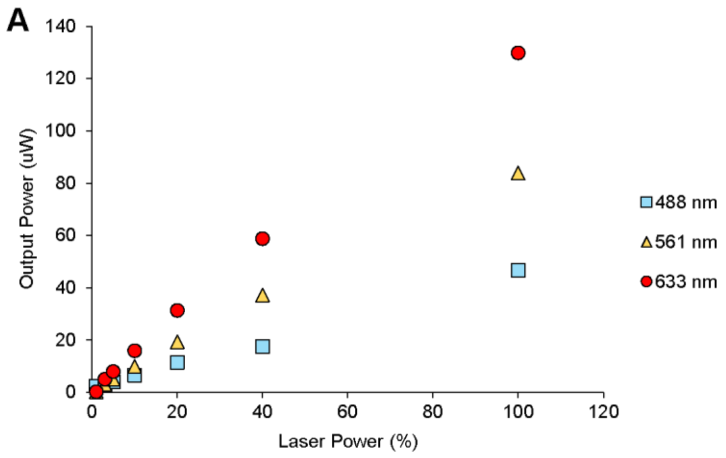


**Figure S1. Relationship between the laser power setting and the light intensity focused through the objective lens.**

(A) Relationship between laser power settings (%) and the light intensity focused through the objective lens (μW) in a Leica SP8 Falcon confocal microscope (DMI8; Leica) equipped with an objective lens, HC PL APO 63×/1.20 W motCORR CS2. Laser power was measured at 488 nm, 531 nm, and 633 nm, which were used in FCCS to illuminate mNG, mScarlet-I/mCherry2, and miRFP670 respectively. The output power was measured by ADCMT 8230E Optical Power Meter.

**Supplementary Figure 2**

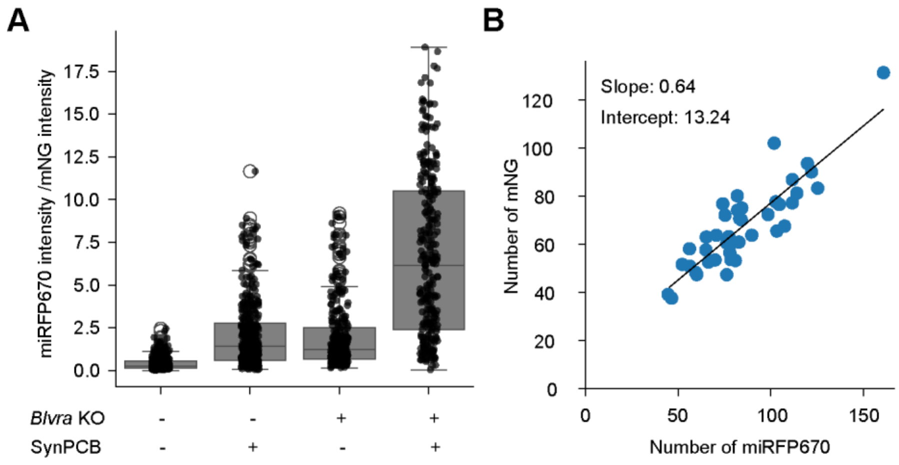


**Figure S2. The effects of *Blvra* KO and SynPCB expression on the fluorescence intensity of miRFP670.**

(A) Normalized fluorescence intensity of miRFP670 in parental or *Blvra* KO HeLa cells expressing mNG-miRFP670 fusion proteins with or without SynPCB. Parental or *Blvra* KO HeLa cells were co-transfected with pCAGGS-mNG-miRFP670 and pCAGGS-SynPCB2.1 (SynPCB+) or pCAGGS-MCS (SynPCB-). The fluorescence intensity of miRFP670 was normalized by that of mNG in the same cell. The box plot of the normalized miRFP670 intensity with the whiskers donating 1.5 times the interquartile range. The box extends from the first to the third quartile. The black dot indicates data from individual cells. (B) Relationship between the number of mNG and that of miRFP670 in cells expressing the mNG-miRFP670 fusion proteins. *Blvra KO* HeLa cells were co-transfected with pCAGGS-mNG-miRFP670 and pCAGGS-SynPCB2.1 (SynPCB+). Cells were analyzed by FCCS to quantify the number of fluorescently labeled mNG and miRFP670. Blue dots indicate data from individual cells and the black line indicates a linear regression line with a slope of 0.63 and a y-intercept of 13.24. These data suggest that approximately 35% of mNG does not fluoresce in HeLa cells.

### Supplementary Figure 3

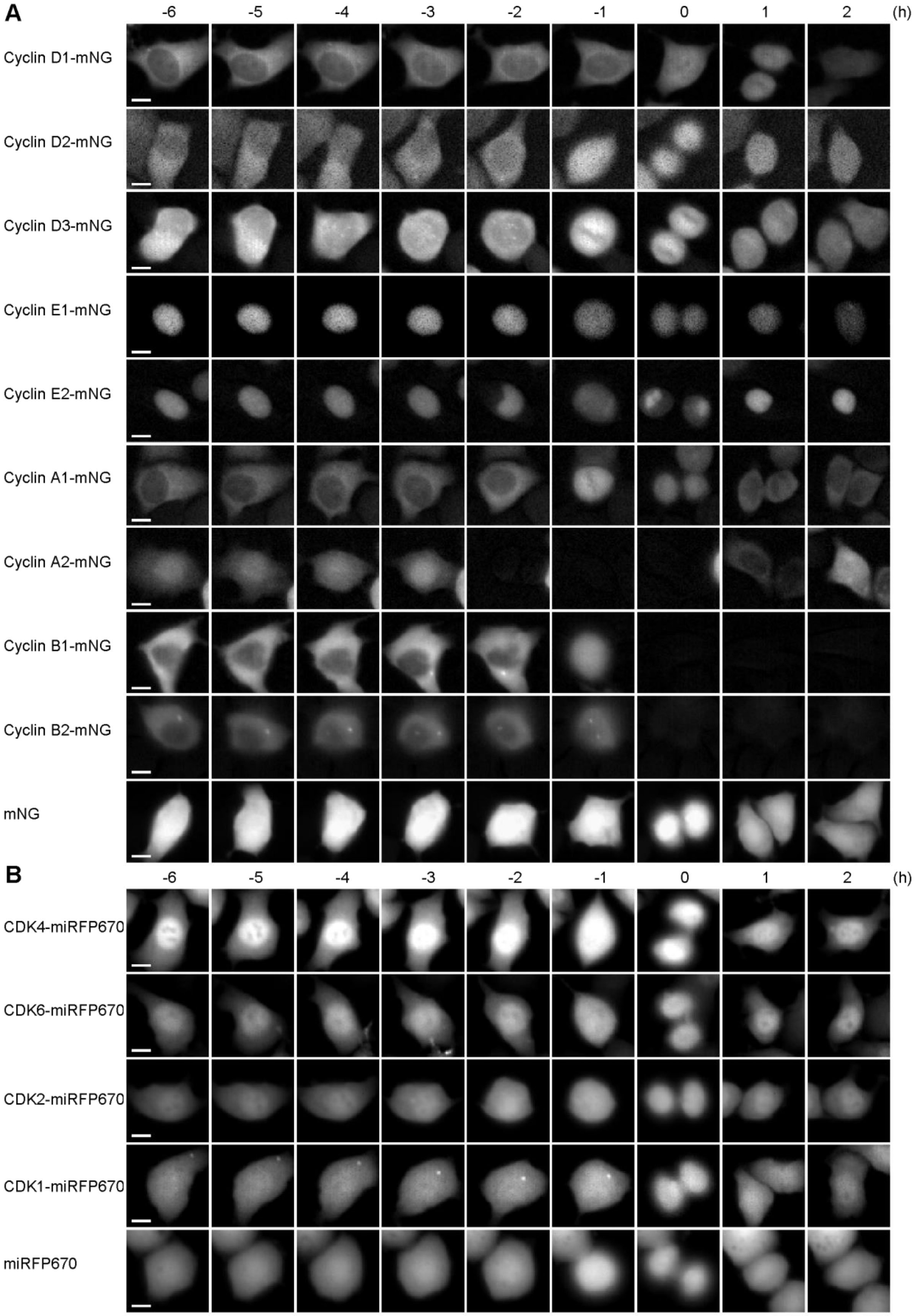

**Figure S3. Localization of cyclin-mNG and CDK-miRFP670 in HeLa cells.**

Subcellular localization of cyclin-mNG (A) and CDK-miRFP670 (B) during the cell cycle in *Blvra* KO HeLa expressing SynPCB. The *Blvra* KO HeLa cells were treated with 1000 ng/uL doxycycline to induce SynPCB expression, and one day after the doxycycline addition, the cells were transfected with pCAGGS-cyclin-mNG or pCAGGS-CDK-miRFP670. 24-48 hours after the transfection, the cells were time-lapse imaged for 24 hours. The representative cells are shown at the indicated time, where time = 0 corresponds to mitosis.

### Supplementary Figure 4

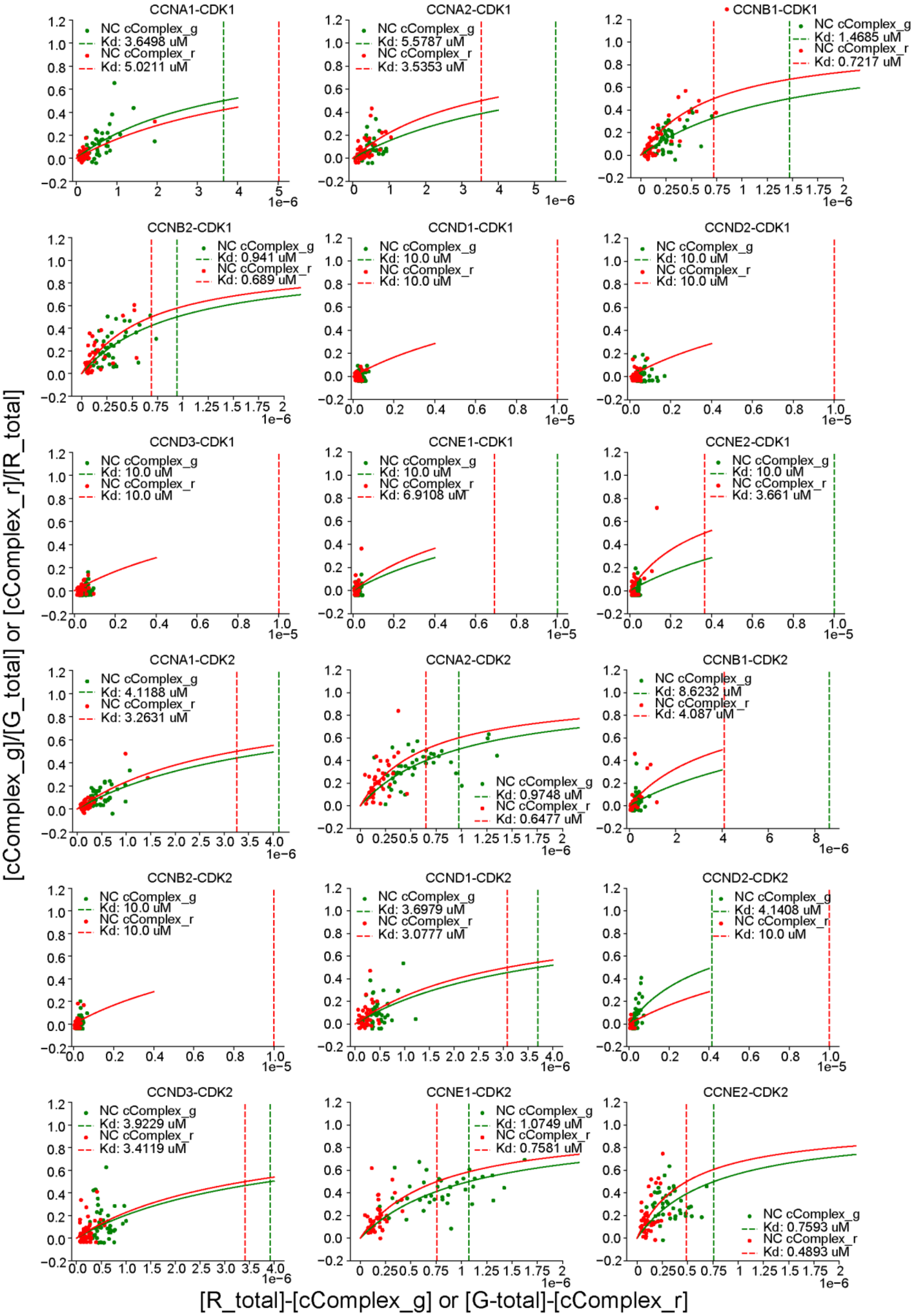

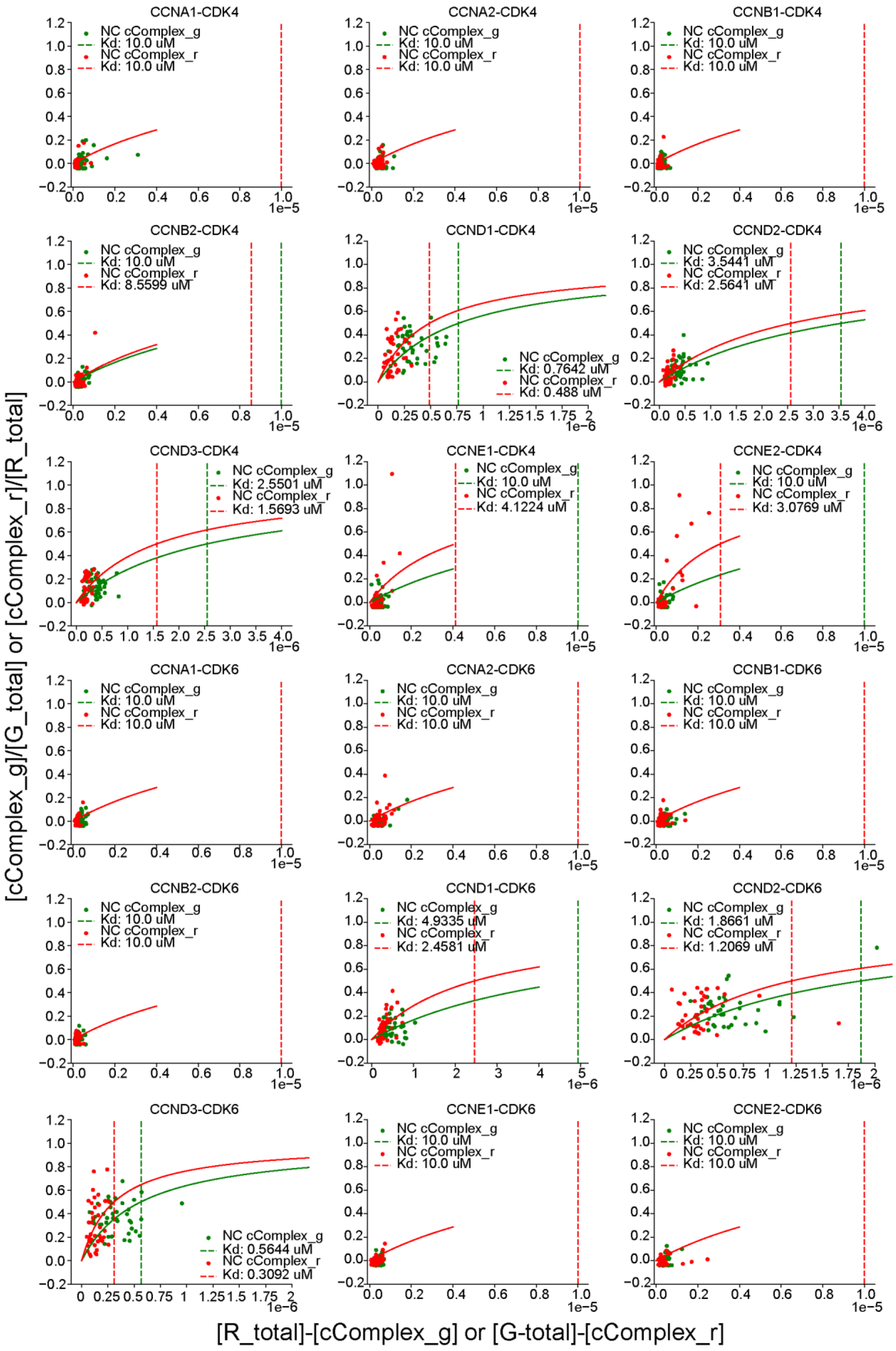

CCNA1-CDK4
NC cComplex_g
Kd: 10.0 uM
NC cComplex_r
Kd: 10.0 uM
CCNA2-CDK4
NC cComplex_g
Kd: 10.0 uM
NC cComplex_r
Kd: 10.0 uM
CCNB1-CDK4
NC cComplex_g
Kd: 10.0 uM
NC cComplex_r
Kd: 10.0 uM
CCNB2-CDK4
NC cComplex_g
Kd: 10.0 uM
NC cComplex_r
Kd: 8.5599 uM
CCND1-CDK4
NC cComplex_g
Kd: 0.7642 uM
NC cComplex_r
Kd: 0.488 uM
CCND2-CDK4
NC cComplex_g
Kd: 3.5441 uM
NC cComplex_r
Kd: 2.5641 uM
CCND3-CDK4
NC cComplex_g
Kd: 2.5501 uM
NC cComplex_r
Kd: 1.5693 uM
CCNE1-CDK4
NC cComplex_g
Kd: 10.0 uM
NC cComplex_r
Kd: 4.1224 uM
CCNE2-CDK4
NC cComplex_g
Kd: 10.0 uM
NC cComplex_r
Kd: 3.0769 uM
CCNA1-CDK6
NC cComplex_g
Kd: 10.0 uM
NC cComplex_r
Kd: 10.0 uM
CCNA2-CDK6
NC cComplex_g
Kd: 10.0 uM
NC cComplex_r
Kd: 10.0 uM
CCNB1-CDK6
NC cComplex_g
Kd: 10.0 uM
NC cComplex_r
Kd: 10.0 uM
CCNB2-CDK6
NC cComplex_g
Kd: 10.0 uM
NC cComplex_r
Kd: 10.0 uM
CCND1-CDK6
NC cComplex_g
Kd: 4.9335 uM
NC cComplex_r
Kd: 2.4581 uM
CCND2-CDK6
NC cComplex_g
Kd: 1.8661 uM
NC cComplex_r
Kd: 1.2069 uM
CCND3-CDK6
NC cComplex_g
Kd: 0.5644 uM
NC cComplex_r
Kd: 0.3092 uM
CCNE1-CDK6
NC cComplex_g
Kd: 10.0 uM
NC cComplex_r
Kd: 10.0 uM
CCNE2-CDK6
NC cComplex_g
Kd: 10.0 uM
NC cComplex_r
Kd: 10.0 uM
1e−5
1e−6
[cComplex_g]/[G_total] or [cComplex_r]/[R_total]
[R_total]-[cComplex_g] or [G-total]-[cComplex_r]

**Figure S4. Fitted curves for calculating the $K_d$ in HeLa cells.**

The data with the fitted curve for calculating the $K_d$ of each cyclin-CDK pair. (Eq. 9) and (Eq. 10) in Material and Method were used for fitting.

**Supplementary Table S1. Plasmid list**

| Plasmid name | Source | Benchling Link |
|---|---|---|
| pCAGGS-mNG | This study | https://benchling.com/s/seq-6pqRQanS6ThPCtiaNBOb?m=slm-X6puXjgySeEjVxZJpqgR |
| pCAGGS-miRFP670 | This study | https://benchling.com/s/seq-upXxqHguFojK71rxty5b?m=slm-aqlU0chqP7alcpdSvc4P |
| pCAGGS-mNG-miRFP670 | This study | https://benchling.com/s/seq-XCnZzrPmXNzuY1dINKxD?m=slm-UJ3vYZktc8OMNI37HH9M |
| pCAGGS-cyclin A1-mNG | This study | https://benchling.com/s/seq-9zWgI1z51hj65j48Y3Gl?m=slm-R0iEDtwa0XnszL4KDd4M |
| pCAGGS-cyclin A2-mNG | This study | https://benchling.com/s/seq-wfUCmsdtBIWvNf1apRRM?m=slm-DHyGVxDP4XgD7HdlAcCZ |
| pCAGGS-cyclin B1-mNG | This study | https://benchling.com/s/seq-MdlhEYv3C6dDM11oAKtc?m=slm-1r7sR1r1suw5Fvdh1AuB |
| pCAGGS-cyclin B2-mNG | This study | https://benchling.com/s/seq-vGLwvpJAF7eJsDiKHYDq?m=slm-xBv57tIDB6VtS3dRnRdr |
| pCAGGS-cyclin D1-mNG | This study | https://benchling.com/s/seq-ExONP5pvsP1i41rbbRdM?m=slm-EhAQMHX4KKJVxlF7DXYs |
| pCAGGS-cyclin D2-mNG | This study | https://benchling.com/s/seq-YZA7PdlOkvoJqfQ0D9Ro?m=slm-mrLi2FWZwKqREr9n3qBJ |
| pCAGGS-cyclin D3-mNG | This study | https://benchling.com/s/seq-Eo9HYVHiNyKthVgvGUe3?m=slm-crcTfy2fdpoMRc9VaCEd |
| pCAGGS-cyclin E1-mNG | This study | https://benchling.com/s/seq-JjO15nirc2pI23oiyKrf?m=slm-bFrx6Ks5BfCvsBkeOvHq |
| pCAGGS-cyclin E2-mNG | This study | https://benchling.com/s/seq-8sra9fUv2Pp3t5DRZXEY?m=slm-szn0AGUXqkkImryxzi4X |
| pCAGGS-synPCB2.1 | Uda et al., 2020 | https://benchling.com/s/seq-371EyyssDZy5KicI0SFX?m=slm-f1armxaOF55aBZcogYqT |

| | | |
|---|---|---|
| pFA6a-mNeonGreen (S.p codon optimized)-kan | Sakai et al.,2021 | https://benchling.com/s/seq-r5vPvd2m4SeMm6VxVhiN |
| pFA6a-miRFP670-hyg | This study | https://benchling.com/s/seq-D2lteVHMldFRlFcrfkF2?m=slm-LH4pwjVuo3FLVCuU2R27 |
| pSKI-BSD-1L-A1-M-SynPCB2.1 | Sakai et al.,2021 | https://benchling.com/s/seq-t5O2P2WzKmrmTYCcEtUk |
| pSKI-NAT-2L-A1-M-spmNeonGreen (S.p codon optimized) | Sakai et al.,2021 | https://benchling.com/s/seq-5KTJy6ykRRzXgNaRUUeV |
| pMKAN3RA1-miRFP670 | This study | https://benchling.com/s/seq-rp2CQhZ5DoGseGE4gJwL?m=slm-AI7G10mqAy7lCWnjICe4 |
| pMNAT2LA1-mCherry2-HA-mNeonGreen | This study | https://benchling.com/s/seq-JLn5XjQsJXGvHHaH2bL5?m=slm-2zOHjXtcZx612NEUvbDz |
| pNATZA1-mScarlet-I-HA-mNeongreen | This study | https://benchling.com/s/seq-4WnrxmvHRSTdw5EAFTGS |
| pMNAT2LA1(ver.2)-miRFP670-spmNeonGreen | This study | https://benchling.com/s/seq-h7wy92qCZupXUuOrREVk?m=slm-UZWIbJwZORvMW9e2IpQe |

**Supplementary Table S2. Fission yeast strain list**

| **Genotype** | **Genotype** | **Genotype** | **Genotype** |
|---|---|---|---|
| h- 1L::Padh1-SynPCB2.1<<bsd 2L::Padh1-miRFP670-spmNeonGreen<<nat | 1C, 1F, 1I, 3C, 3E, 3H | This study | HS490 (LabID) |
| h+ ade6-M210 leu1-32 z::Padh1-mScarlet-I-HA-mNeonGreen<<nat | 1G, 1I | This study | YG181 (LabID) |
| h- 2L::Padh1-mCherry2-HA-mNeonGreen<<nat | 1H, 1I | This study | HS445 (LabID) |
| h- 2L::Padh1-spmNeonGreen<<nat 3R::Padh1-miRFP670<<kan 1L::synPCB2.1<<bsd | 3D, 3E, 3H | This study | HS565 (LabID) |
| h- 1L::Padh1-SynPCB2.1<<bsd cdc2-miRFP670<<hyg cdc13-spmNeonGreen<<kan | 3F, 3H | This study | HS494’ (LabID) |
